# *Paupar* LncRNA Promotes KAP1 Dependent Chromatin Changes And Regulates Subventricular Zone Neurogenesis

**DOI:** 10.1101/187302

**Authors:** Ioanna Pavlaki, Farah Alammari, Bin Sun, Neil Clark, Tamara Sirey, Sheena Lee, Dan J Woodcock, Chris P Ponting, Francis G Szele, Keith W Vance

## Abstract

Many long non-coding RNAs (lncRNAs) are expressed during central nervous system (CNS) development, yet their *in vivo* roles and molecular mechanisms of action remain poorly understood. *Paupar*, a CNS expressed lncRNA, controls neuroblastoma cell growth by binding and modulating the activity of genome-wide transcriptional regulatory elements. We show here that *Paupar* transcript directly binds KAP1, an essential epigenetic regulatory protein, and thereby regulates the expression of shared target genes important for proliferation and neuronal differentiation. *Paupar* promotes KAP1 chromatin occupancy and H3K9me3 deposition at a subset of distal targets, through formation of a DNA binding ribonucleoprotein complex containing *Paupar*, KAP1 and the PAX6 transcription factor. *Paupar*-KAP1 genome-wide co-occupancy reveals a 4-fold enrichment of overlap between *Paupar* and KAP1 bound sequences. Furthermore, both *Paupar* and Kap1 loss of function *in vivo* accelerates lineage progression in the mouse postnatal subventricular zone (SVZ) stem cell niche and disrupts olfactory bulb neurogenesis. These observations provide important conceptual insights into the *trans*-acting modes of lncRNA-mediated epigenetic regulation, the mechanisms of KAP1 genomic recruitment and identify *Paupar* and *Kap1* as regulators of SVZ neurogenesis.

## INTRODUCTION

A subset of nuclear long noncoding RNAs (lncRNAs) have been shown to act as transcription and chromatin regulators using multiple different regulatory mechanisms. These include local functions close to the sites of lncRNA synthesis (Engreitz, Haines et al., 2016) as well as distal modes of action across multiple chromosomes (Chalei, Sansom et al., 2014, Vance, Sansom et al., 2014). Moreover, lncRNA regulatory effects may be mediated by the act of lncRNA transcription as well as RNA sequence dependent interactions with transcription factors and chromatin regulatory proteins (Rutenberg-Schoenberg, Sexton et al., 2016, Vance & Ponting, 2014). Some lncRNAs have been proposed to act as molecular scaffolds to facilitate the formation of multi-component ribonucleoprotein regulatory complexes (Ilik, Quinn et al., 2013, Maenner, Muller et al., 2013, Tsai, Manor et al., 2010, Yang, Flynn et al., 2014, Zhao, Ohsumi et al., 2010), whilst others may act to guide chromatin regulatory complexes to specific binding sites genome wide (Vance & Ponting, 2014). Studies of *cis*-acting lncRNAs such as *Haunt* and *Hottip* have shown that lncRNA transcript accumulation at their sites of expression can effectively recruit regulatory complexes (Pradeepa, McKenna et al., 2017, Yin, Yan et al., 2015). LncRNAs however have also been reported to directly bind and regulate genes across multiple chromosomes away from their sites of synthesis (Carlson, Quinn et al., 2015, Chalei et al., 2014, Chu, Qu et al., 2011, Vance et al., 2014, West, Davis et al., 2014). By way of contrast, the mechanisms by which such *trans*-acting lncRNAs mediate transcription and chromatin regulation at distal bound target genes are less clear.

LncRNAs show a high propensity to be expressed in brain nuclei and cell types relative to other tissues (Mercer, Dinger et al., 2008, Mercer, Qureshi et al., 2010, Ponjavic, Oliver et al., 2009). The adult neurogenic stem cell-containing mouse subventricular zone (SVZ) contributes to brain repair and can be stimulated to limit damage, but is also a source of tumours (Bardella, Al-Dalahmah et al., 2016, Chang, Adorjan et al., 2016). During SVZ lineage progression GFAP+ neural stem cells (NSC) give rise to Mash1+ and Dlx+ transit amplifying progenitors (TAPs) which in turn generate doublecortin+ neuroblasts that migrate to the olfactory bulbs (OB) (Doetsch, Caille et al., 1999). 8,992 lncRNAs are expressed in the SVZ, many of which are differentially expressed during SVZ neurogenesis, suggesting that at least some of these transcripts may play regulatory roles (Ramos, Diaz et al., 2013). However, only a minority of SVZ expressed lncRNAs have been analysed functionally and the full scope of their molecular mechanisms of action remain poorly understood.

*Kap1* encodes an essential chromatin regulatory protein that plays a critical role in embryonic development and in adult tissues. Kap1^-/-^ mice die prior to gastrulation while hypomorphic *Kap1* mouse mutants display multiple abnormal embryonic phenotypes, including defects in the development of the nervous system (Cammas, Mark et al., 2000, Herzog, Wendling et al., 2011, Shibata, Blauvelt et al., 2011). KAP1 interacts with chromatin binding proteins such as HP1 and the SETDB1 histone-lysine N-methyltransferase to control heterochromatin formation and to silence gene expression at euchromatic loci (Iyengar & Farnham, 2011). Despite this fundamental role in epigenetic regulation, the mechanisms of KAP1 genomic targeting are not fully understood. KAP1 does not contain a DNA binding domain but was originally identified through its interaction with members of the KRAB zinc finger (KRAB-ZNF) transcription factor family. Subsequent studies however revealed that KRAB-ZNF interactions cannot account for all KAP1 genomic recruitment events. KAP1 preferentially localises to the 3’ end of zinc finger genes as well as to many promoters and intergenic regions in human neuronal precursor cells. A mutant KAP1 protein however that is unable to interact with KRAB-ZNFs still binds to promoters suggesting functionally distinct subdomains (Iyengar, Ivanov et al., 2011). This work points to the presence of alternative, KRAB-ZNF independent mechanisms that operate to target KAP1 to a distinct set of genomic binding sites. We reasoned that this may involve specific RNA-protein interactions between KAP1 and chromatin bound lncRNAs.

The CNS expressed intergenic lncRNA *Paupar* represents an ideal candidate chromatin-enriched lncRNA with which to further define *trans-*acting mechanisms of lncRNA mediated gene and chromatin regulation. *Paupar* is transcribed upstream from the *Pax6* transcription factor gene and acts to control proliferation and differentiation of N2A neuroblastoma cells *in vitro* (Vance et al., 2014). *Paupar* regulates *Pax6* expression locally, physically associates with PAX6 protein and interacts with distal transcriptional regulatory elements to control gene expression on multiple chromosomes in N2A cells in a dose-dependent manner. Here, we show that *Paupar* directly interacts with KAP1 in N2A cells and that together they control the expression of a shared set of target genes enriched for regulators of neural proliferation and differentiation. Our findings indicate that *Paupar*, KAP1 and PAX6 physically associate on chromatin within the regulatory region of shared target genes and that *Paupar* knockdown reduces both KAP1 chromatin association and histone H3 lysine 9 tri-methylation (H3K9me3) at PAX6 co-bound locations. Genome-wide occupancy maps further identified a preferential enrichment in the overlap between *Paupar* and KAP1 binding sites on chromatin. Our results also show that both *Paupar* and KAP1 loss of function *in vivo* accelerates lineage progression in the mouse postnatal SVZ stem cell niche and disrupts olfactory bulb neurogenesis. We propose that *Paupar* and Kap1 are novel regulators of SVZ neurogenesis, and that *Paupar* operates as a transcriptional cofactor to promote KAP1 dependent chromatin changes at a subset of bound regulatory elements in *trans* via association with non-KRAB-ZNF transcription factors such as PAX6.

## RESULTS

### *Paupar* directly binds the KAP1 chromatin regulatory protein in mouse neural cells in culture

The lncRNA *Paupar* binds transcriptional regulatory elements across multiple chromosomes to control the expression of distal target genes in N2A neuroblastoma cells (Vance et al., 2014). Association with transcription factors such as PAX6 assist in targeting *Paupar* to chromatin sites across the genome. As *Paupar* depletion does not alter PAX6 chromatin occupancy (Vance et al., 2014) we hypothesized that *Paupar* may recruit transcriptional cofactors to PAX6 and other neural transcription factors to regulate gene expression. To test this, we sought to identify transcription and chromatin regulatory proteins that bind both *Paupar* and PAX6 in N2A cells in culture. *In vitro* transcribed biotinylated *Paupar* was therefore immobilised on streptavidin beads and incubated with N2A cell nuclear extract in a pull down assay. Bound proteins were washed, eluted and identified using mass spectrometry (Fig 1a). This identified a set of 78 new candidate *Paupar*-associated proteins that do not bind a control RNA of similar size, including 28 proteins with annotated functions in the control of gene expression that might function as transcriptional cofactors (Fig 1b and Supplemental Table S1).

**Figure 1.**
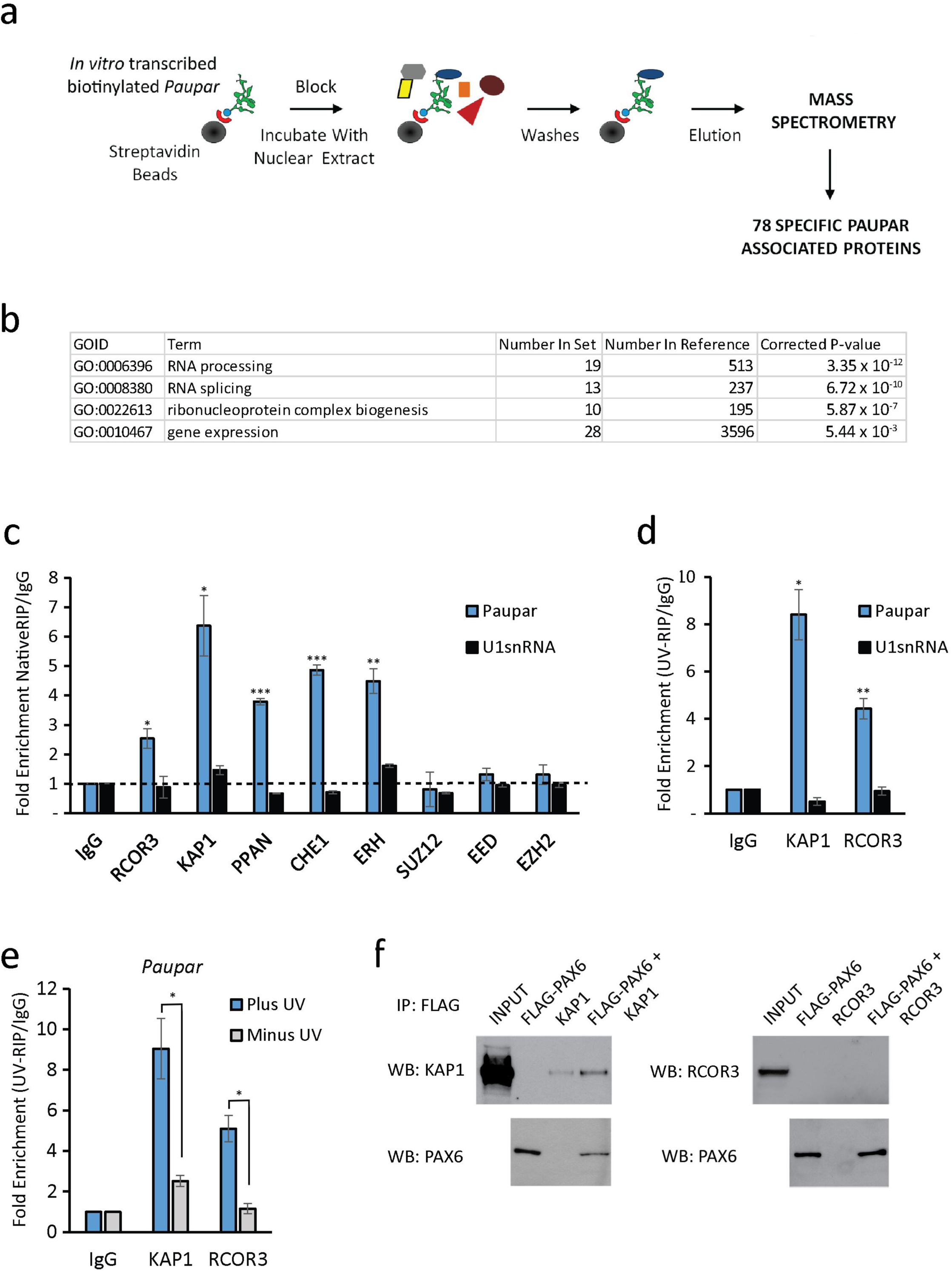
*Paupar* directly binds the KAP1 chromatin regulatory protein in mouse N2A neuroblastoma cells. (a) Overview of the pull down assay. *In vitro* transcribed biotinylated *Paupar* RNA was immobilised on streptavidin beads and incubated with N2A cell nuclear extract. Bound RNA protein complexes were extensively washed and specific *Paupar* associated proteins, which do not interact with a control mRNA of a similar size, identified by mass spectrometry. (b) Gene Ontology terms were used to annotate *Paupar* associated proteins according to biological process. The Bonferroni correction was used to adjust the P-values to correct for multiple testing. (c) Endogenous *Paupar* transcript interacts with transcription and chromatin regulatory proteins in N2A cells. *Paupar* association with the indicated proteins was measured using native RNA-IP. Whole cell lysates were prepared and the indicated regulatory proteins immuno-precipitated using specific antibodies. Bound RNAs were purified and the levels of *Paupar* and the *U1snRNA* control detected in each RIP using qRT-PCR. *Paupar* transcript directly interacts with KAP1 and RCOR3 in N2A cells. Nuclear extracts were prepared from UV cross-linked (d) and untreated (e) cells and immuno-precipitated using either anti-KAP1, anti-RCOR3 or a rabbit IgG control antibody. Associated RNAs were stringently washed and purified. The levels of *Paupar* and a *U1snRNA* control transcript were detected in each UV-RIP using qRT-PCR. Results are presented as fold enrichment relative to control antibody. Mean values +/- SEM., N=3. One-tailed t-test, unequal variance *p<0.05, **p<0.01, ***p<0.001 (f) PAX6 associates with KAP1 in N2A cells. FLAG-PAX6 and KAP1 or RCOR3 expression vectors were transfected into N2A cells. Lysates were prepared two days after transfection and FLAG-PAX6 protein immuno-precipitated using anti-FLAG beads. Co-precipitated proteins were detected by western blotting.

We next performed native RNA-IP experiments in N2A cells to validate potential associations between the endogenous *Paupar* transcript and five gene expression regulators. These were: RCOR3, a member of the CoREST family of proteins that interact with the REST transcription factor; KAP1, a key epigenetic regulator of gene expression and chromatin structure; PPAN, a previously identified regulator of *Pax6* expression in the developing eye; CHE-1, a polymerase II interacting protein that functions to promote cellular proliferation and block apoptosis; and ERH, a transcriptional cofactor that is highly expressed in the eye, brain and spinal cord.

The results revealed that the *Paupar* transcript, but not a non-specific control RNA, was >2-fold enriched using antibodies against RCOR3, KAP1, ERH, PPAN or CHE1 compared to an IgG isotype control in a native RNA-IP experiment (Fig 1c). In addition, *Paupar* did not associate above background with SUZ12, EED and EZH2 Polycomb proteins used as negative controls. This served to further confirm the specificity of the *Paupar* lncRNA-protein interactions because Polycomb proteins associate with a large number of RNAs (Davidovich, Wang et al., 2015) and yet were not identified as *Paupar* interacting proteins in our pull down assay. The endogenous *Paupar* transcript therefore associates with proteins involved in transcription and chromatin regulation in proliferating N2A cells.

To characterise *Paupar* lncRNA-protein interactions further we used UV-RNA-IP to test whether *Paupar* interacts directly with any of these five cofactors. These data showed that *Paupar*, but not an *U1snRNA* control, is highly enriched using antibodies against KAP1 or RCOR3 compared to an IgG control (Fig 1d). A lower level of *Paupar* enrichment is found with CHE1 whereas ERH or PPAN do not appear to interact directly with *Paupar* (Supplemental Fig S1a). Furthermore, the association of *Paupar* with both KAP1 and RCOR3 was reduced in the absence of UV treatment (Fig 1e). These results therefore indicate that the endogenous *Paupar* transcript directly and specifically associates with RCOR3 and KAP1 transcriptional cofactors in neural precursor-like cells in culture.

As a first step to determine whether KAP1 or RCOR3 can act as PAX6 associated transcriptional cofactors we performed immunoprecipitation experiments in N2A cells using transfected FLAG-tagged PAX6 and with HA-KAP1 or HA-RCOR3 proteins. Immunoprecipitation of FLAG-tagged PAX6 using anti-FLAG beads co-immunoprecipitated transfected KAP1 protein, but not RCOR3 (Fig 1f), suggesting that PAX6 and KAP1 are present within the same multi-component regulatory complex. Consistent with this, a previous study showed that KAP1 interacts with PAX3 through the amino terminal paired domain, which is structurally similar in PAX6, to mediate PAX3 dependent transcriptional repression (Hsieh, Yao et al., 2006). Together, these results indicate that KAP1 may regulate *Paupar* and PAX6 mediated gene expression programmes.

### *Paupar* and KAP1 control expression of a shared set of target genes that are enriched for regulators of neuronal function and cell cycle in N2A cells

KAP1 regulates the expression of genes involved in the self-renewal and differentiation of multiple cell types, including neuronal cells (Iyengar & Farnham, 2011) and thus is an excellent candidate interactor for mediating the transcriptional regulatory function of *Paupar*. To investigate whether *Paupar* and KAP1 functionally interact to control gene expression we first tested whether they regulate a common set of target genes. We depleted *Kap1* expression in N2A cells using shRNA transfection and achieved approximately 90% reduction in both protein (Fig 2a) and transcript (Fig 2b) levels. *Paupar* levels do not change upon KAP1 knockdown indicating that KAP1 dependent changes in gene expression are not due to regulation of *Paupar* expression (Fig 2b). Transcriptome profiling using microarrays then identified 1,913 differentially expressed genes whose expression significantly changed (at a 5% false discovery rate [FDR]) greater than 1.4-fold (log2 fold change ≈ 0.5) upon KAP1 depletion (Fig 2c and Supplemental Table S2). 282 of these genes were previously identified to be regulated by human KAP1 in Ntera2 undifferentiated human neural progenitor cells (Iyengar et al., 2011). Transient reduction of *Kap1* expression by approximately 55% using a second shRNA expression vector (*Kap1* shB) also induced expression changes for 7 out of 8 *Kap1* target genes with known functions in neuronal cells that were identified in the microarray (Supplemental Fig 1b). These data further validate the specificity of the KAP1 regulated gene set.

**Figure 2.**
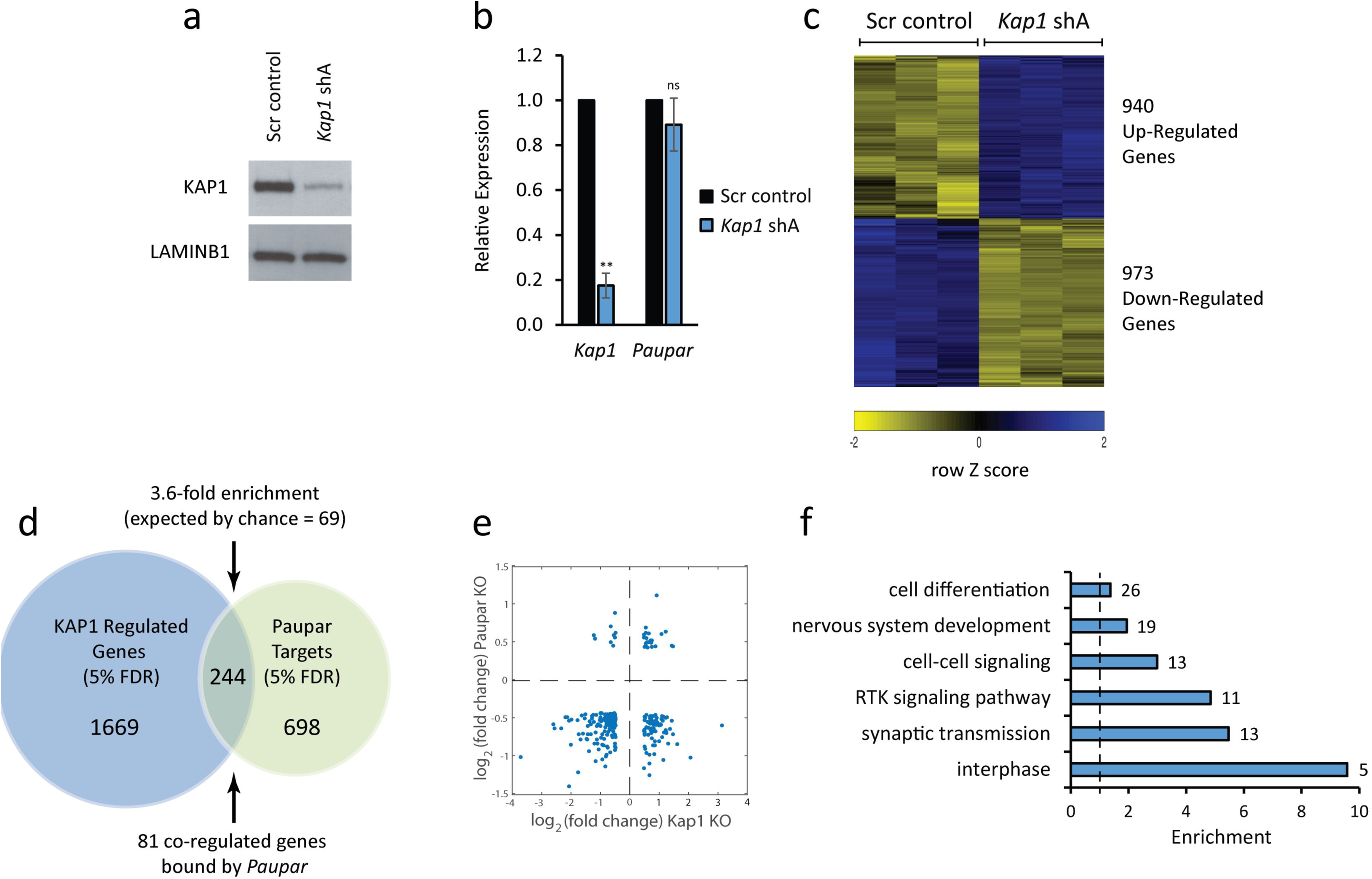
*Paupar* and KAP1 regulate shared target genes involved in neural cell proliferation and function. N2A cells were transfected with either the shA *Kap1* targeting shRNA expression vector or a scrambled control and pTK-Hyg selection plasmid. Three days later cells were expanded and hygromycin was added to the medium to remove untransfected cells. (a) After seven days, western blotting was performed to determine KAP1 protein levels. LAMINB1 was used as a loading control. (b) *Kap1* and *Paupar* transcript levels were analysed by qRT-PCR. Data was normalised using *Gapdh* and expression changes are shown relative to a non-targeting scrambled control (set at 1). Mean values +/- SEM., N=3. One-tailed t-test, unequal variance **p<0.01 (c) KAP1 regulated genes were identified using a GeneChip Mouse Gene 1.0 ST Array (5% FDR, log2 fold change > 0.5). (d) Intersection of *Kap1* and *Paupar* regulated genes revealed common target genes whose expression is controlled by both these factors. (e) The majority (87%) of *Paupar* and *Kap1* shared target genes are positively regulated by *Paupar*. (f) Gene Ontology analysis of *Paupar* and *Kap1* common target genes was performed using GOToolBox. Representative significantly enriched categories were selected from a hypergeometric test with a Benjamini-Hochberg corrected P-value threshold of 0.05.

We previously showed that *Paupar* knockdown induces changes in the expression of 942 genes in N2A cells (Vance et al., 2014). Examination of the intersection of KAP1 and *Paupar* transcriptional targets identified 244 genes whose levels are affected by both *Paupar* and KAP1 knockdown in this cell type (Fig 2d and Supplemental Table S3). This represents a significant 3.6-fold enrichment over the number expected by random sampling and is not due to co-regulation because *Kap1* is not a *Paupar* target (Vance et al., 2014). A large majority (87%; 212/244) of these common targets are positively regulated by *Paupar* and for two-thirds of these genes (161/244) their expression changes in the same direction upon *Paupar* or KAP1 knockdown (Fig 2e). Furthermore, Gene Ontology enrichment analysis of these 244 genes showed that *Paupar* and KAP1 both regulate a shared set of target genes enriched for regulators of interphase, components of receptor tyrosine kinase signalling pathways as well as genes involved in nervous system development and essential neuronal cell functions such as synaptic transmission (Fig 2f). Genes targeted by both *Paupar* and KAP1 are thus expected to contribute to the control of neural stem-cell self-renewal and neural differentiation.

### *Paupar*, KAP1 and PAX6 associate on chromatin within the regulatory region of shared target genes

In order to investigate *Paupar* mediated mechanisms of distal gene regulation we next sought to determine whether *Paupar*, KAP1 and PAX6 can form a ternary complex on chromatin within the regulatory regions of their shared target genes. To do this, we first integrated our analysis of PAX6 regulated gene expression programmes in N2A cells (Vance et al., 2014) and identified 87 of the 244 *Paupar* and KAP1 common targets, which is 35.8-fold greater than expected by random sampling, whose expression is also controlled by PAX6 (Fig 3a and Supplemental Table S3). We found that 34 of these genes contain a CHART-Seq mapped *Paupar* binding site within their GREAT defined putative regulatory regions (Vance, 2016, Vance et al., 2014) and predicted that these represent functional *Paupar* binding events within close genomic proximity to direct transcriptional target genes (Fig 3a and Supplemental Table S3).

**Figure 3.**
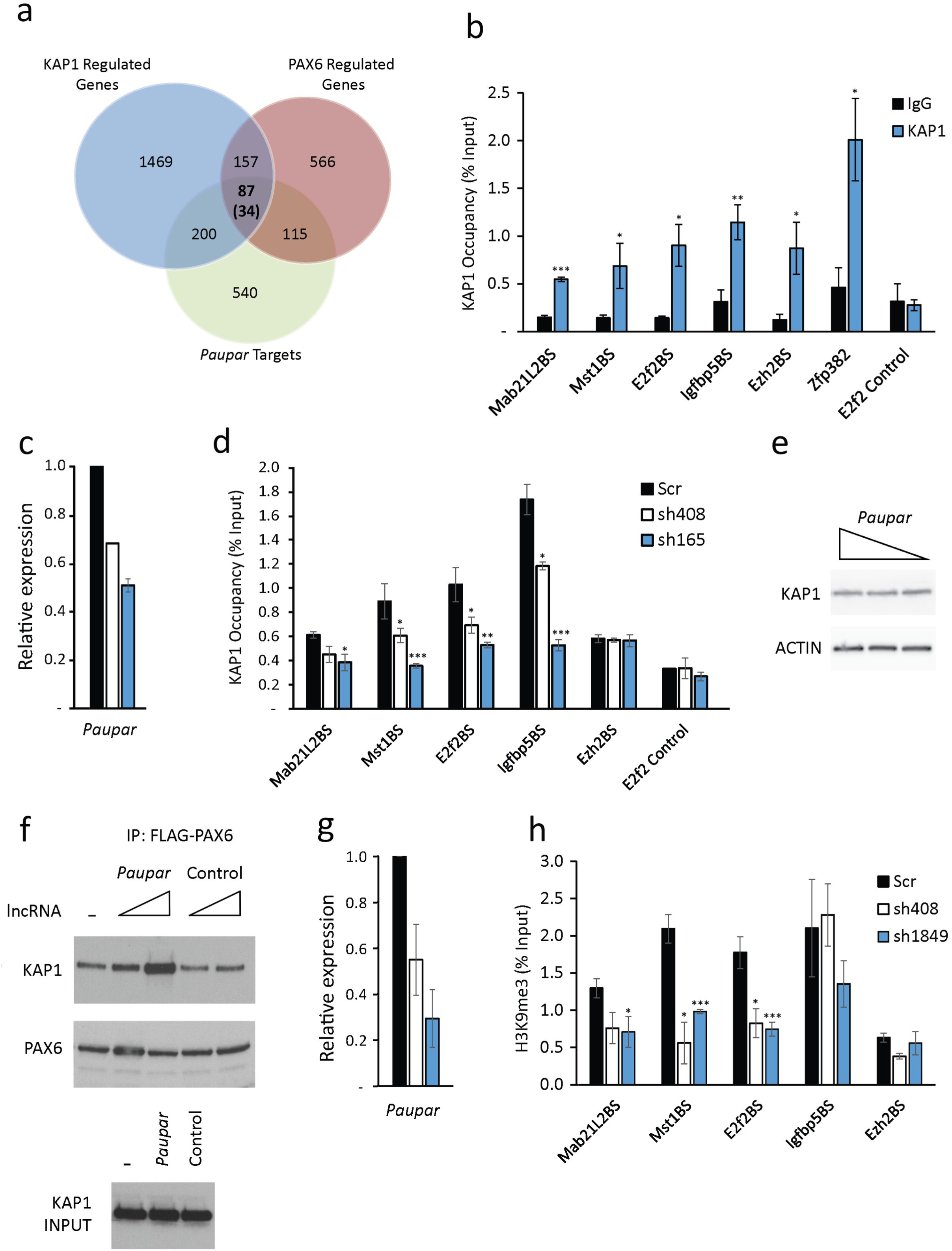
*Paupar* promotes KAP1 chromatin occupancy and H3K9me3 deposition at PAX6 bound sequences within the regulatory regions of common targets. (a) Intersection of *Paupar*, KAP1 and PAX6 regulated genes identified 87 common target genes. 34 of these genes (in brackets) contain a *Paupar* binding site within their regulatory regions. (b) ChIP assays were performed in N2A cells using either an antibody against KAP1 or an isotype specific control. (c) N2A cells were transfected with either a non-targeting control or two independent *Paupar* targeting shRNA expression vectors. Cells were harvested for ChIP three days later. *Paupar* depletion was confirmed using qRT-PCR. (d) *Paupar* knockdown reduces KAP1 chromatin occupancy at shared binding sites. ChIP assays were performed using either an anti-KAP1 polyclonal antibody or a normal IgG rabbit control. (e) Western blotting showed that KAP1 proteins levels do not change upon *Paupar* knockdown. ACTIN was used as a control. (f) *Paupar* promotes PAX6-KAP1 association. FLAG-PAX6 and KAP1 expression vectors were co-transfected into N2A cells along with increasing concentrations of *Paupar* or a size matched control lncRNA expression vector. Expression of the maximum concentration of either *Paupar* or control RNA in each IP does not alter KAP1 input protein levels (lower panel). Lysates were prepared two days after transfection and FLAG-PAX6 protein immuno-precipitated using anti-FLAG beads. The amount of DNA transfected was made equal in each IP using empty vector and proteins in each complex were detected by western blotting. (g, h) *Paupar* knockdown reduces H3K9me3 at a subset of bound sequences in *trans*. For ChIP assays, the indicated DNA fragments were amplified using qPCR. % input was calculated as 100*2^(Ct Input-Ct IP). Results are presented as mean values +/-SEM, N=3. One-tailed t-test, unequal variance *p<0.05, **p<0.01, ***p<0.001

ChIP-qPCR analysis previously identified four of these *Paupar* bound locations within the regulatory regions of the *Mab21L2*, *Mst1*, *E2f2* and *Igfbp5* genes that are also bound by PAX6 in N2A cells (Vance et al., 2014). We therefore measured KAP1 chromatin occupancy at these regions as well as at a negative control sequence within the first intron of *E2f2* using ChIP and identified a specific enrichment of KAP1 chromatin association at the *Mab21L2*, *Mst1*, *E2f2* and *Igfbp5* genes compared to an IgG isotype control (Fig 3b). KAP1 binding to these regions is only 2- to 4-fold reduced compared to the Zfp382 3’ UTR positive control (Fig. 3b) which represents an exemplar high affinity KAP1 binding site (Iyengar et al., 2011). Furthermore, KAP1 and *Paupar* also co-occupy a binding site within the *Ezh2* gene. As *Ezh2* is regulated by *Paupar* and KAP1 but not by PAX6 this suggests that *Paupar* and KAP1 can also interact with specific sites on chromatin using additional PAX6 independent mechanisms. Together, these data suggest that *Mab21L2*, *Mst1*, *E2f2* and *Igfbp5* are co-ordinately regulated by a ribonucleoprotein complex containing *Paupar*-KAP1-PAX6.

### *Paupar* functions as a transcriptional cofactor to promote KAP1 chromatin occupancy and H3K9me3 deposition at PAX6 bound sequences

KAP1 is recruited to its target sites within 3’UTRs of ZNF genes through the association with KRAB-ZNF transcription factors (Iyengar et al., 2011, O’Geen, Squazzo et al., 2007). However, *Paupar* bound sequences are preferentially located at gene promoters and are not enriched for KRAB-ZNF transcription factor binding motifs (Vance et al., 2014). This suggests that *Paupar* may play a role in recruiting KAP1 to a separate class of binding site in a KRAB-ZNF independent manner. To test this, *Paupar* expression was first depleted using transient transfection of *Paupar* targeting shRNA expression vectors (Fig 3c). We then performed ChIP-qPCR to measure KAP1 chromatin occupancy in control and *Paupar* knockdown N2A cells at the four *Paupar*-KAP1-PAX6 co-occupied binding sites within the regulatory regions of the *Mab21L2*, *Mst1*, *E2f2* and *Igfbp5* genes, a *Paupar*-KAP1 bound sequence within the *Ezh2* gene that is not regulated by PAX6, and a control sequence that is not bound by *Paupar*. The results show that KAP1 chromatin binding is significantly decreased at the four *Paupar*-KAP1-PAX6 bound regions upon *Paupar* depletion and that the extent of KAP1 chromatin association appears to be dependent on *Paupar* transcript levels (Fig. 3d). KAP1 chromatin association is also not reduced at the *Ezh2* gene *Paupar*-KAP1 binding site or at the control sequence that is not bound by *Paupar* (Fig 3d), whilst total KAP1 protein levels do not detectably change upon *Paupar* knockdown (Fig 3e), further confirming specificity.

These results imply that *Paupar* functions to promote KAP1 chromatin association at a subset of its genomic binding sites in *trans* and that this requires the formation of a DNA bound ternary complex containing *Paupar*, KAP1 and PAX6. Consistent with this, co-expression of the *Paupar* lncRNA promotes KAP1-PAX6 association in a dose dependent manner in an immunoprecipitation experiment (Fig 3f). This effect is specific for the *Paupar* transcript because expression of a size-matched control RNA does not alter KAP1-PAX6 association. *Paupar* thus functions as a transcriptional cofactor to promote the assembly of a *Paupar*-KAP1-PAX6 ternary complex on chromatin in *trans*. This ribonucleoprotein complex appears to function as a regulator of genes involved in controlling neural stem cell self-renewal and differentiation.

We next tested whether *Paupar* can induce histone modification changes at bound target genes on different chromosomes away from its sites of synthesis. As KAP1 interacts with the SETDB1 methyltransferase to mediate histone H3K9me3 deposition (Schultz, Ayyanathan et al., 2002), we first determined the levels of H3K9me3 at *Mab21L2*, *Mst1*, *E2f2* and *Igfbp5* bound sequences using ChIP-qPCR. This revealed an enrichment of H3K9me3 modified chromatin at all five locations (Supplemental Fig S2), consistent with a previous study showing that many KAP1 bound promoters are marked by H3K9me3 (O’Geen et al., 2007). shRNA mediated reduction of *Paupar* transcript levels using two different shRNAs resulted in a significant decrease in histone H3K9me3 modification at 3 of 4 of these shared binding sites tested using ChIP (Fig. 3g, h). No change in histone H3K9me3 was detected at *Ezh2* gene whose expression does not change upon PAX6 depletion. Together, these data show that *Paupar* functions to modulate KAP1 chromatin association and histone H3K9me3 deposition at a subset of its shared binding sites in *trans*.

### *Paupar* co-occupies an enriched subset of KAP1 binding sites genome-wide

We next examined the intersection between *Paupar* and KAP1 bound locations genome-wide in order to generate a more comprehensive view of the potential of *Paupar* for regulating KAP1 function. ChIP-seq profiling of KAP1 chromatin occupancy showed that KAP1 associates with 5510 genomic locations compared to input DNA in N2A cells (1% FDR) (Supplemental Table S4). KAP1 binding sites are particularly enriched at promoter regions, over gene bodies and at the 3’UTRs of zinc finger genes (Fig. 4a), consistent with previous studies mapping human KAP1 genomic occupancy (Iyengar et al., 2011, O’Geen et al., 2007). Intersection of KAP1 bound locations with our CHART-seq map of *Paupar* genomic binding in N2A cells (Vance et al., 2014) identified 46 KAP1 binding sites that are co-occupied by *Paupar* and not bound in a LacZ negative control CHART-seq pull down (Fig. 4b), only one of which is located within the 3’UTR of a ZNF gene (zfp68) (Supplemental Table S4). Notably, this represents a significant (p < 0.001) 4-fold enrichment of *Paupar* and KAP1 co-occupied locations as estimated using Genome Association Tester (GAT) (Fig. 4b). In addition, plotting the distribution of peak intensities across these co-occupied regions revealed a precise coincidence of *Paupar* and KAP1 binding (Fig. 4c). These data therefore show that *Paupar* co-occupies an enriched subset of KAP1 bound sequences genome-wide and suggest that *Paupar* mediated genomic recruitment of KAP1 may involve interactions with other transcription factors in addition to KRAB-ZNF association.

**Figure 4.**
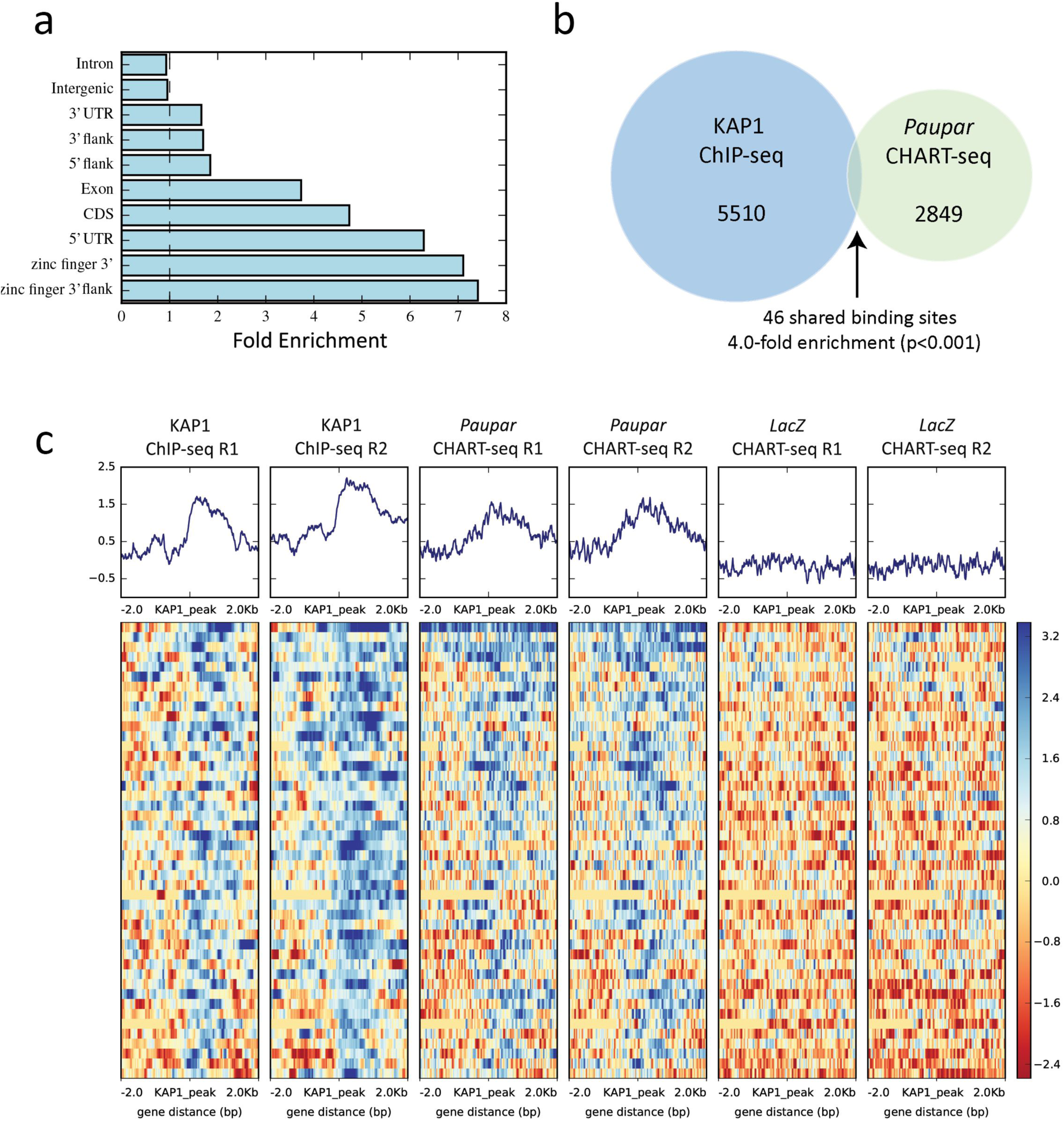
*Paupar* co-occupies a subset of KAP1 binding sites on chromatin genome-wide. 5510 KAP1 binding sites common to both replicates were identified relative to input DNA (1% FDR) (Supplemental Table S4). (a) GAT analysis shows that the sites of KAP1 occupancy are particularly enriched at promoter regions (5’UTRs), over gene bodies and over the 3’UTR exons of zinc finger genes (q = 0.00002). (b) Intersection of KAP1 and *Paupar* binding sites in N2A cells identified 46 KAP1 bound locations that are specifically co-occupied by *Paupar*. This represents a significant (p < 0.001) 4-fold enrichment as estimated using GAT. (c) Sequencing read density distribution over the 46 shared binding locations was calculated and revealed a coincidence of *Paupar* and KAP1 binding site centrality.

### *Paupar* and Kap1 regulate the SVZ neurogenic niche and olfactory bulb neurogenesis

Our results indicate that *Paupar* and KAP1 regulate the expression of shared target genes important for proliferation and neuronal differentiation in N2A cells. We next expanded this observation and tested whether *Paupar* and Kap1 can regulate the same neurodevelopmental process *in vivo*. To do this, we used the mouse SVZ system as it is experimentally convenient for discovering many different neurodevelopmental mechanisms. Lineage progression can be monitored by electroporating the neonatal SVZ; in 24 hrs the NSC are labelled, 3 days post electroporation (3dpe) TAPs appear and by 7 dpe labelled neuroblasts are seen migrating into the OB (Boutin, Diestel et al., 2008a, Chesler, Le Pichon et al., 2008).

We first showed using RT-qPCR that *Paupar* is expressed in the SVZ, as well as in neurospheres cultured from P4 SVZ (Supplemental Fig S3a), and then confirmed the efficiency of the *Paupar* targeting shRNA expression vectors to deplete *Paupar* transcript in neurospheres cultured from P4 SVZ (Supplemental Fig S3b). sh165 caused robust *Paupar* knockdown (KD) whereas sh408 moderately reduced *Paupar* expression enabling us to identify dose dependent regulatory effects. Nucleofection of *Paupar* KD constructs and a scrambled (scr) control plasmid targeted ~60% of cells, as measured using GFP, but we determined *Paupar* levels in all cells. Thus on a cell-by-cell basis the relative level of knockdown is predicted to be greater than shown (Supplemental Fig S3b). To study *Paupar* function in neurogenesis, we electroporated P1 pups with *Paupar* KD constructs or scr controls and examined the SVZ 24 hours post electroporation (24hpe) and 3 days post electroporation (3dpe). To control for differences in the number of cells electroporated in the different groups we measured the percentage of GFP+ cells expressing lineage markers (Fig. 5a, c). Immunostaining showed that at 24hpe, the percent of GFP+ cells expressing the TAP marker MASH1 was increased by more than 50% with sh165 knockdown (Fig. 5b). This was confirmed by immunostaining with the TAP and neuroblast marker DLX2 which showed a greater than 30% increase with both knockdown constructs (Fig. 5b). Additionally we showed that the percentage of GFP+ cells positive for the proliferation marker Ki67 was significantly increased in the sh165 group (Fig. 5b). At 24hpe the majority of cells in scramble controls are radial glia-like neural stem cells (Boutin et al., 2008a, Chesler et al., 2008). These results thus suggest that after *Paupar* KD a larger percentage of cells are progressing into the next phase of the SVZ lineage and are actively proliferating. We next carried out immunohistochemistry for the same markers at 3dpe and quantification showed that fewer GFP+ cells expressed the radial glial/neural stem cell marker GFAP upon KD with the sh165 construct (Fig. 5c-e). This further suggests that *Paupar* loss increases lineage progression and/or diminishes SVZ stem cell maintenance.

**Figure 5.**
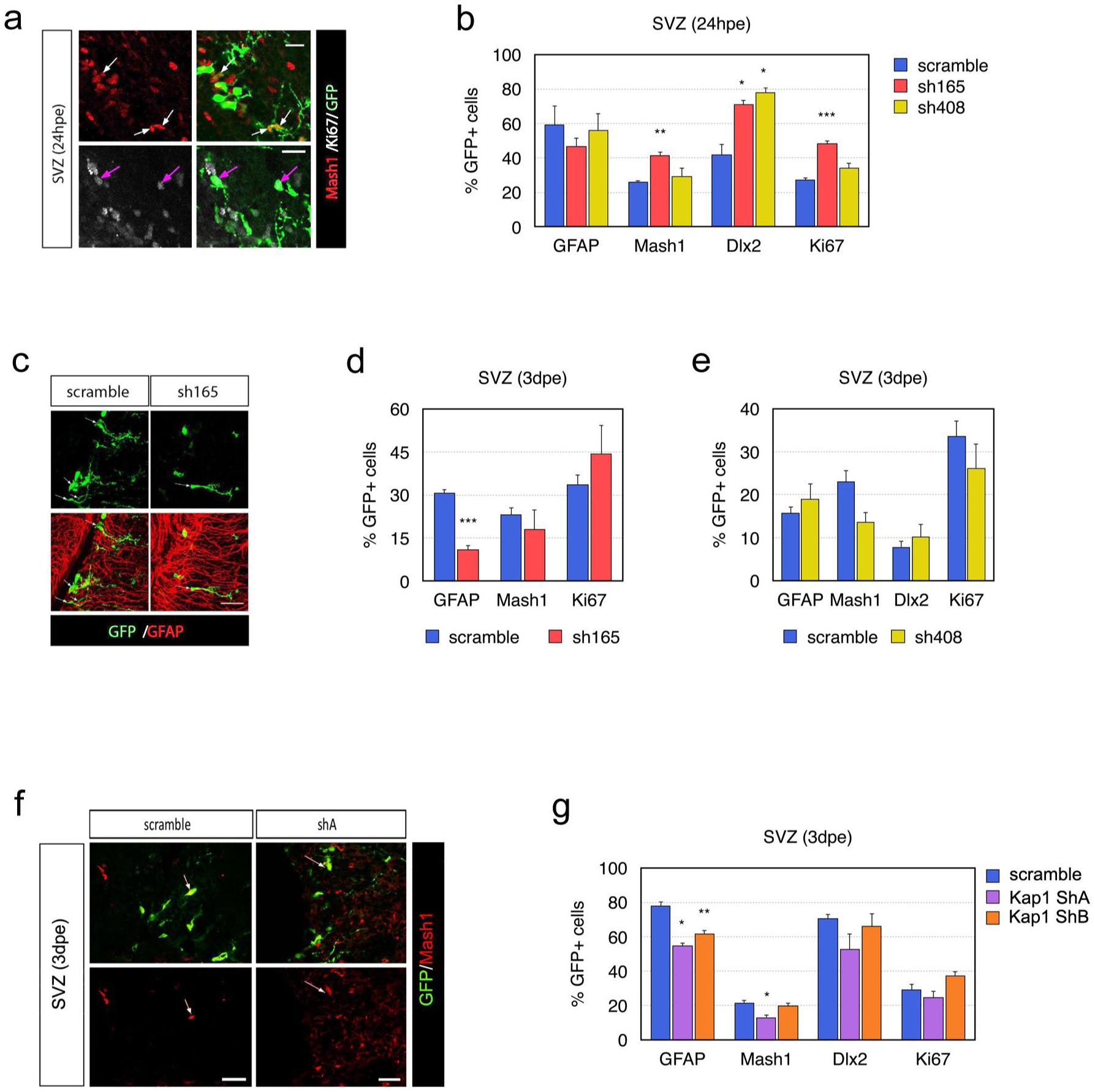
*Paupar* and Kap1 regulate SVZ neurogenesis *in vivo*. P1 pups were electroporated with the indicated shRNA expression vectors. All shRNA plasmids also express GFP. (a) Example of co-immunostaining of MASH1, KI67 and GFP in the SVZ with electroporated GFP+ cells. White arrows in top row indicate MASH1+/GFP+ cells 24hpe. Magenta arrows in bottom row (different field) indicate KI67+/GFP+ cells. (b) Quantification after *Paupar* knockdown of the percent of GFP+ cells in the SVZ that express GFAP, MASH1, DLX2 or KI67 at 24hpe. N≥3. (c) Immunostaining of GFP and GFAP in the SVZ, 3dpe. The small arrows indicate GFAP+/GFP+ cells. (d-e) Quantification after *Paupar* knockdown of the percent of GFP+ cells that express GFAP, MASH1 or KI67 at 3dpe. N≥4. (f) Example of co-immunostaining of GFP and MASH1 in the SVZ, 3dpe. The small arrows indicate MASH1+/GFP+ cells. (g) Quantification after *Kap1* knockdown of the percent of GFP+ cells that express GFAP, MASH1, DLX2 or KI67 at 3 dpe. N=3. Data are shown as mean ± SEM and analysed by two-tailed Student t-tests. *p<0.05, **p<0.01, ***p<0.001. Scale bars represent 20 μm (a), 50 μm (c).

The Allen Brain Atlas shows *Kap1* expression in the SVZ. To study the functional effect of *Kap1* on SVZ neurogenesis, P1 pups were electroporated with either a scr control or the *Kap1* shRNA expression vectors that we used to deplete *Paupar* in N2A cells (Fig. 3 and Supplemental Fig S1b) and sections were immunostained for GFP and SVZ markers (Fig. 5f). At 3dpe of *Kap1* shA and shB, the percentage of GFP+ cells that expressed the radial glial/neural stem cell marker GFAP significantly decreased (Fig. 5g). This is similar to *Paupar* KD and is consistent with accelerated lineage progression. Also similar to *Paupar* KD at 3dpe, *Kap1* KD did not alter the percent of GFP+ cells which expressed DLX2 or Ki67. However, the percentage of MASH1+ cells decreased slightly but significantly at 3dpe post *Kap1* shA KD, which was not found upon *Paupar* KD. Since we showed that *Paupar* and *Kap1* regulate similar as well as different genes this result may be due to differential gene regulation. Furthermore, these *Paupar* and *Kap1* mediated changes in cell subtype numbers are not due to altered rates of cell death because we did not detect changes in the number of CASPASE3+ cells (Supplemental Fig S4a), or in the percentage of GFP+ cells that are Tunel+ between scr control and any of the *Paupar* or *Kap1* shRNA expression vectors (Supplemental Fig S4b, c).

We next studied how *Paupar* or *Kap1* affects the number of electroporated cells that reach the OB 7dpe. There were significantly fewer GFP+ cells in the OB after *Paupar* KD using sh165 KD compared to the scr control whilst KD with sh408 caused a slight but statistically non-significant decrease in OB GFP+ cell numbers (Fig. 6a, b). Co-staining with the immature neuroblast marker DCX (Yang, Sundholm-Peters et al., 2004) showed that all GFP+ cells in the OB were DCX+ and this was not altered by *Paupar* KD (Supplemental Fig S3d). Similar to *Paupar*, at 7dpe of either Kap1 KD construct, there was a significant reduction in the number of GFP+ cells that had migrated from the SVZ to the OB (Fig. 6c, d). These results suggest that both *Paupar* and *Kap1* are required for the production of newborn OB neurons.

**Figure 6.**
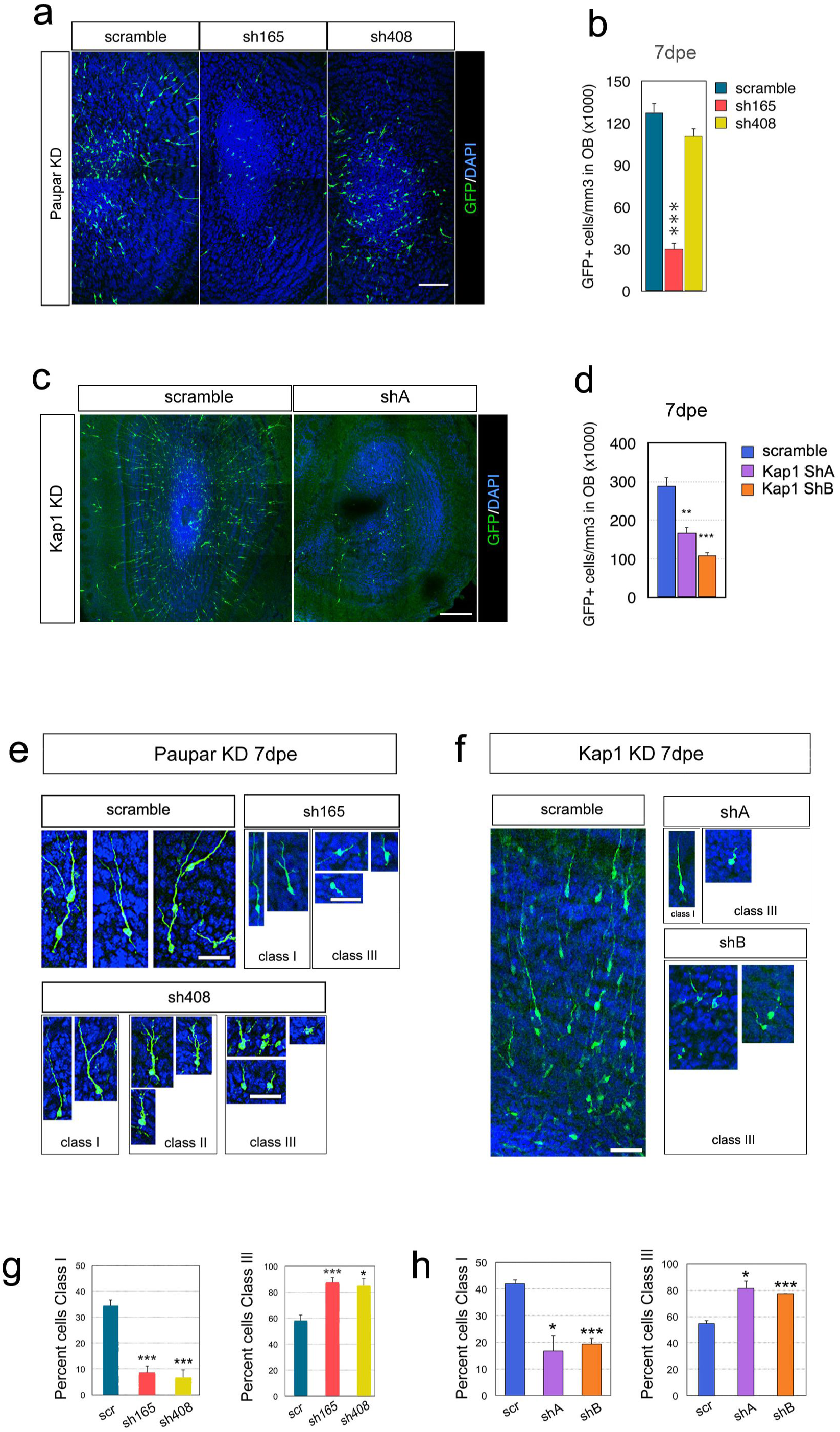
*Paupar* and *Kap1* loss of function alters OB neuron morphology. P1 pups were electroporated with the indicated shRNA expression vectors. All shRNA plasmids also express GFP. (a, b) Immunostaining and quantification of GFP+ cells that were electroporated in the SVZ and migrated to the OB, *Paupar* KD, 7dpe. N≥3. (c-d) GFP+ cells that have migrated to the olfactory bulb 7dpe decrease after *Kap1* KD. Quantification of the density of electroporated cells in the OB after *Kap1* KD. N=3. (e) High magnification showing different morphologies in GFP+ granule layer OB neurons 7 dpe, *Paupar* KD. For ease of comparison neuronal orientations were aligned to vertical. The cells shown in the scr control group are class I N≥3. (f) High magnification showing different morphologies in GFP+ granule layer OB neurons 7 dpe, *Kap1* KD. Neuronal orientations rendered vertical. The scr control image shows several class I as well as class III neurons. (N=3). (g) Quantification of the percent of cells with Class I and Class III morphology 7 days after *Paupar* KD. (h) Quantification of the percent of cells with Class I and Class III morphology 7 days after *Kap1* KD. Data are shown as mean ± SEM and analysed by two-tailed Student t-tests. *p<0.05, **p<0.01, ***p<0.001. Scale bars represent 100 μm (a), 200 μm (c), 30 μm (e), 50 μm (f).

Interestingly, *Paupar* as well as *Kap1* knockdown altered the morphology of newborn neurons that migrated to the OB (Fig. 6e-h). In scr controls many GFP+ neurons in the OB granule layer had processes extending radially towards the pial surface and some of the processes were branched and these were classified as class I cells (Fig 6e, f). By contrast, after *Paupar* KD, a variety of abnormal morphologies were observed, which we classified as class II or class III (Fig 6e). Class II cells were rare but were distinguished by many short branched processes. Class III cells were stunted with only short or no processes (Fig 6e). Quantification revealed that after *Paupar* KD the percentage of cells with Class I morphology was 34±2% in scr controls but only 8±3% in sh165 and 6±3% in sh408 (P=0.0005 and P=0.0009, respectively) (Fig. 6g). Conversely, after *Paupar* KD there were more class III neurons in the sh165 group 87±4% as well as in the sh408 group 85±6% compared to 58±5% controls (P=0.003 and P=0.02, respectively). *Kap1* knockdown showed similar effects (Fig 6f, h); shA and shB resulted in 16.7±5.6% and 19.3±2.0% of Class I neurons versus 42.0±1.5% in controls (P=0.012 and P=0.013, respectively). Again, the number of Class III neurons increased from 54.7±2.2% in controls to 81.3±5.6% after shA KD and 77.3±0.3% after shB KD (P=0.0009 and P=0.0005, respectively). These data further suggest that *Kap1* and *Paupar* affect postnatal neurogenesis by disrupting both migration into the OB and the morphology of newborn neurons.

## DISCUSSION

LncRNAs can bind and regulate target genes on multiple chromosomes away from their sites of transcription. Furthermore, the number of lncRNAs that function in this way is steadily increasing suggesting that nuclear lncRNAs are likely to exert a wide range of currently uncharacterised, *trans*-acting functions in transcription and chromatin regulation. Moreover, loss-of-function studies using animal model systems are needed to identify and characterise lncRNA regulatory roles during embryonic development and in adult tissue homeostasis to clarify the importance of this class of transcript *in vivo*.

To gain novel insights into lncRNA gene regulation we investigated the mode of action of the CNS expressed lncRNA *Paupar* at chromosomal binding sites away from its site of synthesis in N2A cells. We show that *Paupar* directly binds the KAP1 epigenetic regulatory protein and thereby regulates the expression of shared target genes important for proliferation and neuronal differentiation. Our data indicate that *Paupar* modulates histone H3K9me deposition at a subset of distal bound transcriptional regulatory elements through its association with KAP1, including at a binding site upstream of the *E2f2* gene. These chromatin changes are consistent with our previous report that this *E2f2* bound sequence functions as a transcriptional enhancer whose activity is restricted by *Paupar* transcript levels (Vance et al., 2014). Our results therefore suggest a model in which *Paupar* directed histone modification changes in *trans* alter the activity of bound regulatory elements in a dose dependent manner.

Several other lncRNAs have also been shown to alter the chromatin structure of target genes in *trans*. These include the human *PAUPAR* orthologue which can inhibit H3K4 tri-methylation of the *Hes1* promoter in eye cancer cell lines, as well as lncRNA-HIT which induces p100/CBP mediated changes in histone H3K27ac at bound sequences to regulate genes involved in chondrogenesis (Carlson et al., 2015, Ding, Wang et al., 2016). The lncRNA *Hotair* is one of the most studied *trans*-acting lncRNAs. Whilst *Hotair* has been proposed to guide PRC2 to specific locations in the genome to induce H3K27me3 and silence gene expression (Chu et al., 2011), recent conflicting studies report that PRC2 associates with low specificity to lncRNAs and suggest that *HOTAIR* does not directly recruit PRC2 to the genome to silence gene transcription (Davidovich et al., 2015, Kaneko, Son et al., 2013, Portoso, Ragazzini et al., 2017). Mechanistic studies on individual *trans*-acting lncRNAs such as *Paupar* are therefore needed to further define general principles of genome-wide lncRNA transcription and chromatin regulation.

It is proposed that lncRNAs may guide chromatin modifying complexes to distal regions in the genome though RNA-RNA associations at transcribed loci, or either directly through RNA-DNA base pairing or indirectly through RNA-protein-DNA associations (Rutenberg-Schoenberg et al., 2016, Vance & Ponting, 2014). We show that *Paupar* acts to increase KAP1 chromatin association by promoting the formation of a DNA binding regulatory complex containing *Paupar*, KAP1 and PAX6 within the regulatory regions of shared target genes in *trans*, as illustrated in the model in Fig 7. This suggests that *Paupar* functions as a cofactor for transcription factors such as PAX6 to modulate target gene expression across multiple chromosomes. In a similar manner, *Prncr1* and *Pcgem1* lncRNAs interact with the androgen receptor (AR) and associate with non-DNA binding cofactors to facilitate AR mediated gene regulation (Yang, Lin et al., 2013). LncRNA mediated recruitment of chromatin regulatory proteins to DNA bound transcription factors may represent a common mechanism of *trans*-acting lncRNA gene regulation, in line with their suggested role as molecular scaffolds (Tsai et al., 2010).

**Figure 7.**
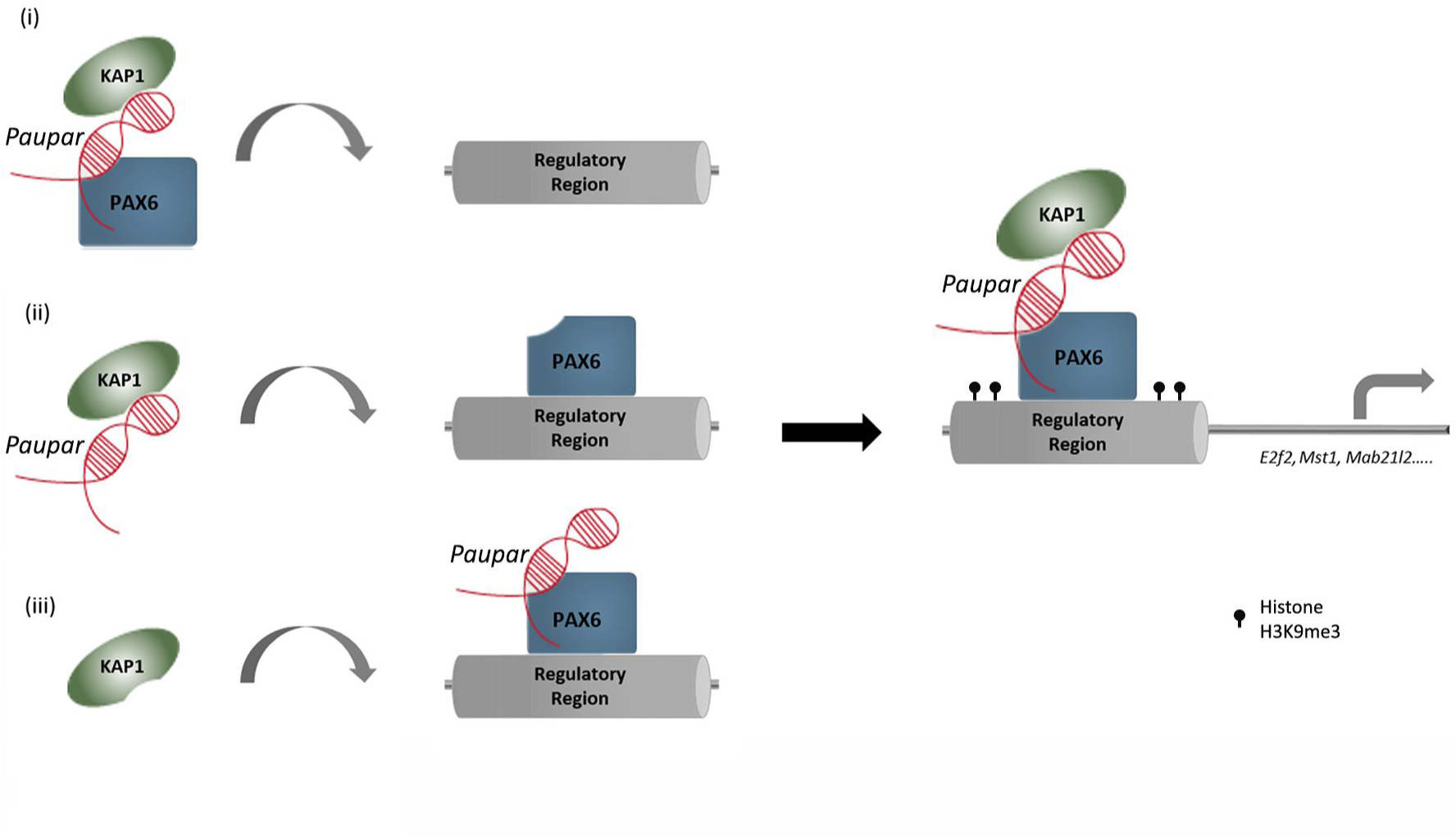
Schematic detailing possible *Paupar* mode of action at distal bound regulatory regions. *Paupar* promotes KAP1 chromatin association and H3K9me3 deposition through the assembly of a DNA bound ribonucleoprotein complex containing *Paupar*, KAP1 and PAX6 within the regulatory regions of the *Mab21L2*, *Mst1*, *E2f2* and *Igfbp5* direct target genes. We propose three potential (non-mutually exclusive) scenarios to describe the order of assembly of this complex: (i) A ternary complex forms in the nucleoplasm before binding DNA; (ii) *Paupar* interacts with KAP1 and guides it to DNA bound PAX6; or (iii) KAP1 is recruited to a DNA bound *PAX6-Paupar* complex. This leads to local H3K9me3 modification changes at these bound sequences in *trans*. The model was generated taking into consideration the discovery that *Paupar* genome wide binding sites contain an enrichment of motifs for neural transcription factors but are not enriched for sequences that are complementary to *Paupar* itself (Vance et al., 2014). This suggests that *Paupar* does not bind DNA directly but is targeted to chromatin indirectly through RNA-protein interactions with transcription factors such as PAX6. Moreover, KAP1 is a non-DNA binding chromatin regulator that is also targeted to the genome through interactions with transcription factors.

KAP1 is guided to 3’UTR of zinc finger genes in the genome through association with KRAB-ZNF transcription factors (O’Geen et al., 2007). However, the mechanisms of KAP1 genome-wide recruitment are not fully understood (Iyengar et al., 2011). Our data identify KAP1 as a novel RNA binding protein and show that *Paupar* plays a role in modulating the recruitment of KAP1 to specific PAX6 bound locations in the genome. We further assessed the extent to which *Paupar* may be able to modulate KAP1 genome-wide recruitment and identified 46 shared binding sites on chromatin, only one of which was within a 3’ UTR of a zinc finger gene. These results raise the possibility that additional chromatin enriched lncRNAs may operate to recruit KAP1 to specific locations in the genome and that this may involve context specific interactions with both KRAB-ZNF as well as non KRAB-ZNF containing transcription factors such as PAX6.

Our knockdown studies indicate that *Paupar* and *Kap1* are required for normal postnatal SVZ neurogenesis *in vivo*. Neonatal SVZ stem cells are the cells lining the ventricles postnatally and are thus are initially targeted by electroporation with TAPs appearing after 3 days and neuroblasts after one week (Boutin et al., 2008a, Chesler et al., 2008). Reduced *Paupar* expression increased proliferation in stem cells suggesting it normally maintains stem cell quiescence and restricts lineage progression. Supporting this, the TAP markers MASH1 and DLX2 increased one day after *Paupar* KD. Importantly, *Mash1* is necessary for stem cell activation (Andersen, Urban et al., 2014) and maintaining neurogenic proliferation (Castro, Martynoga et al., 2011). Similarly, *Dlx2* is necessary for SVZ neurogenesis (Brill, Snapyan et al., 2008) and stimulates lineage progression (Suh, Obernier et al., 2009). Therefore, increased MASH1 and DLX2 levels after *Paupar* KD likely accelerate lineage progression. *Gfap* expression is precipitously lost as neonatal SVZ stem cells transition to TAPS (Doetsch, Garcia-Verdugo et al., 1997). Three days after *Paupar* knockdown GFAP decreased, further suggesting that *Paupar* negatively regulates lineage progression. Similarly, *Kap1* knockdown decreased *Gfap* expression, suggesting *Paupar* and *Kap1* may have other SVZ functions in common. However, *Kap1* but not *Paupar* KD decreased MASH1 levels 3dpe possibly due to the fact that they regulate common as well as distinct programmes of gene expression. Both *Paupar* and *Kap1* loss-of-function reduced the number newborn neurons in the OB. Accelerated lineage progression does not predict reduced OB neurogenesis and the SVZ effects may not be directly linked the OB effects. We controlled for apoptosis and showed that neither *Paupar* nor *Kap1* seems to regulate apoptosis in the SVZ neurogenic system. However, fewer newborn neuroblasts had healthy morphology and more had stunted morphology after *Paupar* or *Kap1* knockdown.

This study identifies *Paupar* and *Kap1* as novel regulators of SVZ neurogenesis *in vivo* and provides important conceptual insights into the distal modes of lncRNA mediated gene regulation. Given the widespread role played by *Kap1* in genome regulation and chromatin organisation we anticipate that further chromatin associated lncRNAs will be found to functionally interact with KAP1.

## MATERIALS AND METHODS

### Plasmid Construction

*Kap1* targeting short hairpin RNAs (shRNAs), designed using the Whitehead Institute siRNA selection program, were synthesized as double stranded DNA oligonucleotides and ligated into pBS-U6-CMVeGFP as shown previously (Vance et al., 2014). The *Paupar* targeting sh165 and sh408 expression constructs, the non-targeting scrambled control shRNA and pCAGGS-*Paupar* expression vector are also detailed in (Vance et al., 2014). To generate the PAX6 expression vector, *Pax6* coding sequence was PCR amplified from mouse N2A cell cDNA as a NotI-XhoI fragment and inserted into pcDNA3.1(+) (Invitrogen). The forward primer incorporated a DNA sequence to insert the DYKDDDDK FLAG epitope tag in frame at the amino terminal end of PAX6. *Rcor3* coding sequence was also PCR amplified from mouse N2A cell cDNA and cloned into pcDNA3.1(+) to generate pcDNA-RCOR3. pcDNA3-HA-KAP1 was a kind gift from Colin Goding (Ludwig Institute, Oxford). The sequences of the oligonucleotides used in this study are listed in Supplemental Table S5.

### Cell Culture

N2A mouse neuroblastoma cells (ATCC CCL-131) were grown in DMEM supplemented with 10% foetal bovine serum. All transfections were performed using FuGENE 6 (Promega) following the manufacturer’s instructions. To generate *Kap1* knockdown cells, approximately 2 × 10^5^ cells were plated per well in a six well plate. 16–24 h later cells were transfected with 1.5 μg *Kap1* shRNA expression construct and 300 ng (5:1 ratio) pTK-Hyg (Clontech). Three days after transfection, cells were trypsinised, resuspended in growth medium containing 200 μg/ml Hygromycin B and plated onto a 6 cm dish. Drug resistant cells were grown for 7 days and harvested as a pool.

### Immunoprecipitation

1 x 10^6^ N2A cells were seeded per 10 cm dish. The next day, cells were transfected with different combinations of pcDNA3-FLAG-PAX6, pcDNA3-Myc-KAP1, pCAGGS-Paupar, pCAGGS-AK034351 control transcript or pcDNA3.1 empty vector. 6 μg plasmid DNA was transfected in total. Two days later, cells were washed twice with ice-cold PBS, transferred to 1.5 ml microcentrifuge tubes and lysed in 1 ml ice-cold IP Buffer (IPB) (50 mM Hepes pH 7.5, 350 mM NaCl, 1 mM MgCl_2_, 0.5 mM EDTA and 0.4% IGEPAL CA-630) for 30 min, 4°C with rotation. Lysates were pelleted at 14,000 rpm, 20 min, 4°C in a microfuge, supernatant was added to 30 μl anti-FLAG M2 Magnetic Beads (#M8823, Sigma) and incubated overnight at 4°C with rotation. Beads were washed three times with IPB and eluted in 20 μl Laemmli sample buffer for 5 min at 95°C. Bound proteins were detected by Western Blotting using anti-FLAG M2 (F3165, Sigma), anti-KAP1 (ab10483, Abcam), anti-RCOR3 (A301-273A, Bethyl Laboratories) and Protein A HRP (ab7456, Abcam).

### RNA Pull Down Assay

Sense RNA was *in vitro* transcribed from pCR4-TOPO-*Paupar* using T7 RNA polymerase, according to manufacturer’s instructions (New England Biolabs). Transcribed RNA was concentrated and purified using the RNeasy MinElute Cleanup kit (Qiagen). Purified RNA was then 5’ end labelled with biotin-maleimide using a 5’ EndTag nucleic acid labelling system (Vector laboratories). Streptavidin coated Dynabeads M-280 (Invitrogen) were washed, prepared for RNA manipulation and the 5’ biotinylated RNA bound according to manufacturer’s instructions. N2A cell nuclear extract was diluted in affinity binding/washing buffer (150 mM NaCl, 50 mM HEPES, pH 8.0, 0.5% Igepal, 10 mM MgCl_2_) in the presence of 100ug/ml tRNA, 40U/ml RNaseOUT (Invitrogen) and a protease inhibitor cocktail (Roche). RNA coated beads were incubated with nuclear extract at room temperature for 2 hours with rotation. The supernatant was then removed, the beads washed six times (10 min) with affinity/binding washing buffer, and bound protein eluted by heating to 95°C in the presence of Laemmli sample buffer for 5 min. Samples were loaded onto a 10% Tris-glycine polyacrylamide gel (BioRad) and subjected to denaturing SDS-PAGE until they just entered the resolving gel. Protein samples were then excised, diced, and washed three times with nanopure water. Tryptic digest and mass spectrometry were performed by the Central Proteomics Facility (Dunn School of Pathology, University of Oxford).

### RNA-IP

Approximately 1x10^7^ N2A cells were used per RNA-IP. Native RNA-IP experiments were performed using the Magna RIP Kit (Millipore) according to the manufacturer’s instructions. UV-RIP was carried out as described in (Vance et al., 2014). We used the following rabbit polyclonal antibodies: anti-RCOR3 (A301-273A, Bethyl Laboratories), anti-CoREST (07-455, Millipore), anti-KAP1 (ab10483, Abcam), anti-ERH (ab96130, Abcam), anti-PPAN (11006-1-AP, Proteintech Group) and rabbit IgG (PP64B, Millipore).

### Chromatin Immunoprecipitation

4 x 10^6^ N2A cells per ChIP were seeded in 15 cm plates. The next day, cells were transfected with either 15 μg *Paupar* targeting shRNA expression vectors or a non-targeting scr control. Three days later cells were harvested for ChIP using either 5 μg anti-KAP1 (ab10483, Abcam), anti-histone H3K9me3 (39161, Active Motif) or normal rabbit control IgG (#2729, Cell Signalling Technology) antibodies. ChIP was performed as described in (Vance et al., 2014). For KAP1 ChIP-seq the following modifications were made to the protocol: approximately 2x10^7^ N2A cells per ChIP were double cross-linked, first using 2 mM disuccinimidyl glutarate (DSG) for 45 min at room temperature, followed by 1% formaldehyde for 15 min at room temperature, as described in (Nowak, Tian et al., 2005). Chromatin was sheared to approximately 200 bp using a Bioruptor Pico (Diagenode) and ChIP DNA and matched input DNA from two independent KAP1 ChIP experiments were sequenced on an Illumina HiSeq 4000 (150 bp paired-end sequencing).

### ChIP-seq Analysis

The Babraham Bioinformatics *fastqscreen* (https://www.bioinformatics.babraham.ac.uk/projects/fastq_screen/) and *fastQC*(https://www.bioinformatics.babraham.ac.uk/projects/fastqc/) tools were used to screen the raw reads for containments and to assess quality. We removed traces of the adapter sequence from the raw reads using the *Trimmomatic* tool (Bolger, Lohse et al., 2014). Trimmomatic was also used to trim by quality with the options: *LEADING:3 TRAILING:3 SLIDINGWINDOW:4:15 MINLEN:50*. The trimmed reads were aligned to the mm10 reference genome, using the Burrows-Wheeler Aligner (Li & Durbin, 2010) with the command: > *bwa mem mm10 <pair_1.fq> <pair_2.fq*>. Alignment quality was assessed with the *Qualimap 2.2.1* tool (Okonechnikov, Conesa et al., 2016). The aligned reads were filtered to exclude reads with a MAPQ alignment quality <20. Furthermore, we excluded reads aligning to blacklisted regions identified by the ENCODE consortium (Consortium, 2012). MACS2 version 2.1.1.20160309 was used to identify genomic regions bound by KAP1. We further filtered the aligned reads to retain only those with length 150 and called peaks relative to the input controls using the options ‘ --*gsize=1.87e9–qvalue=0.01-B–keep-dup auto*’. To examine the read density distribution in the vicinity of KAP1 peaks we used *deepTools* (Ramirez, Ryan et al., 2016). Read density was calculated with respect to input using the bamCompare tool from deepTools, with the option ‘--*binSize 10*’. The matrix of read densities in the vicinity of KAP1 peaks was calculated using ‘*computeMatrix reference-point*’, and heatmaps plotted with ‘*plotHeatmap*’. The Genomic Association Test tool *GAT* (Heger, Webber et al., 2013) was used to characterise KAP1 binding sites and the relationship between KAP1 and *Paupar*. Coordinates with respect to the mm10 reference genome for characteristic genomic regions (exons, introns, 3’ UTRs, etc) were downloaded from the UCSC Genome Table Browser (https://genome.ucsc.edu/cgi-bin/hgTables). The enrichment of KAP1 peaks and the intersection of KAP1 and *Paupar* peaks with respect to these genomic regions was assessed using GAT with the options ‘–ignore-segment-track–num-samples=100000’ and using the complement of the blacklist regions as the workspace. To test for significance coincidence of KAP1 and *Paupar* peaks we use GAT with the same options. The *Paupar* CHART-Seq peakset from (Vance et al., 2014) was used for comparison.

### Transcriptomic Analysis

Total RNA was isolated from triplicate control and KAP1 knockdown cells using the Qiagen Mini RNeasy kit following the manufacturer’s instructions. RNA samples with a RNA Integrity Number greater than 8, as assessed on a BioAnalyzer (Agilent Technologies), were hybridised to Mouse Gene 1.0 ST Arrays as detailed in (Chalei et al., 2014). Differentially expressed genes were identified and Gene Ontology analysis was performed as previously (Vance et al., 2014).

### Neurosphere Assay

Neurospheres were cultured according to standard protocols as previously described (Dizon, Shin et al., 2006). In brief, age P3-P6 CD1 pups were anesthetized by hypothermia and decapitated, and the brains were immediately dissected out and sectioned in the coronal plane with a McIlwain tissue chopper. The SVZ was then dissected out in ice-cold HBSS in a sterile laminar flow hood. Accutase was used for 15 mins for dissociation. Cells were cultured in defined Neurobasal media supplemented with 20ng/ml EGF (Sigma) and 20ng/ml bFGF (R&D). Cells were seeded at a density of 100 cells/μl and passaged every 3-4 days.

### Neural stem cell nucleofection

3-4 x 10^6^ dissociated neurosphere cells were nucleofected according to the protocol of LONZA (VPG-1004). Cells were mixed with 100μl nucleofection solution (82μl of Nucleofector Solution + 18μl of supplement) and 5- 10μg DNA and transferred into cuvettes. 500μl of culture medium was added into the cuvette and the sample was then transferred into 1ml medium and centrifuged at 1200rpm for 5min and resuspended with fresh medium and plated at 200 000 cells/ 2ml in a polyheme coated 6-well plate.

### Postnatal electroporation

Electroporation was performed as published (Boutin, Diestel et al., 2008b, Chesler et al., 2008). DNA plasmids were prepared with Endofree Maxi kit (Qiagen) and mixed with 0.1% fast green for tracing. DNA concentrations were matched in every individual experiment. P1 CD1 pups were anesthetized with hypothermia and 1-2 μl of plasmids were injected with glass capillary. Electrical pulses (100V, 50ms ON with 850ms intervals for 5 cycles) were given with tweezer electrodes (CUY650P5). Pups were recovered, then returned to dam and analysed at the indicated time.

### Immunohistochemistry and imaging

Immunohistochemistry was as previously described (Young, Al-Dalahmah et al., 2014). The following primary antibodies were used: mouse anti-MASH1 (1:100, BD Pharmingen), rabbit anti-KI67 (1:500, Abcam), rabbit anti-CASPASE3 (1:1000, Cell Signaling), rabbit anti-mCherry (1:500, Abcam 167453), rat anti-GFAP (1:500, Invitrogen), chicken anti-GFP (1:500, Aves), rabbit anti-DLX2 (1:50, Abcam). The secondary antibodies were Alexafluor conjugated (Invitrogen). In situ cell death detection kit (Tunel), TMR red (cat# 12156 792910) was used to detect apoptosis. Sections were imaged with Zeiss 710 Laser Scanning Microscopy. For co-localization in GFP+ cells, a 40X oil immersion objective was used and 2μm intervals were used for generating Z-stacks. Confocal images were analysed with ImageJ.

### Morphological evaluation

All GFP+ neuroblasts in the granule layer of the OB were binned into Class I, II, or III groups. Only cells with obvious cell bodies and that were entirely found in the field were included. Cells in the rostral migratory stream in the core of the OB, and in OB layers outside of the granule layer were not included. N=3-5 mice per group.

### Ethics

All mouse experiments were performed in accordance with institutional and national guidelines and regulations under UK Home Office Project Licence PPL 3003311.

### Data availability

Microarray and ChIP-Seq data will be deposited in the GEO database.

## ACKNOWLEDGEMENTS

This project has been funded by a Biotechnology and Biological Sciences Research Council grant to KWV (BB/N005856/1; KWV, IP), a Medical Research Council (MR/M010554/1; FGS, BS, FA) grant to FS, and the European Research Council (Project Reference 249869, DARCGENs), the Medical Research Council and Wellcome Trust (CPP, TS).

## SUPPLEMENTAL MATERIAL

**Supplemental Tables**

Supplemental Table S1: Specific *Paupar* Associated Proteins

Supplemental Table S2: KAP1 Regulated Genes

Supplemental Table S3: Shared Target Genes

Supplemental Table S4: KAP1 ChIP-seq binding locations

Supplemental Table S5: Oligos

**Supplemental Figure Legends**

**Supplemental Figure S1.**
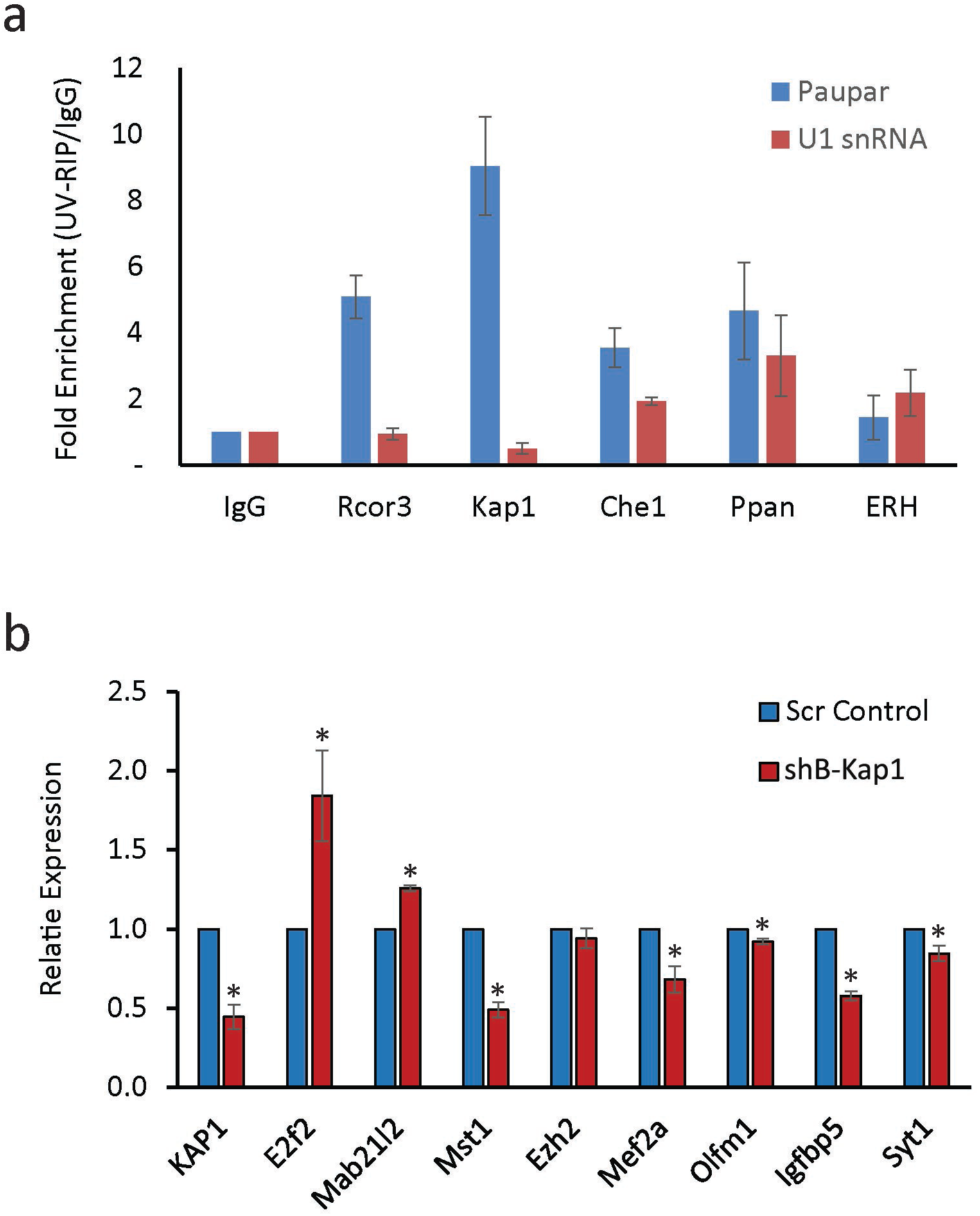
(a) Nuclear extracts were prepared from UV cross-linked N2A cells and immuno-precipitated using either the indicated antibodies or a rabbit IgG control antibody. Associated RNAs were stringently washed and purified. The levels of *Paupar* and *U1snRNA* were detected in each UV-RIP using qRT-PCR. Results are presented as fold enrichment relative to control antibody. Mean values +/- SEM. (b) N2A cells were transfected with an additional Kap1 targeting shRNA expression vector shB-Kap1 or a scrambled control plasmid. Three days later cells were harvested and expression analysed using RT-qPCR. Samples were normalised using *Gapdh* and the results are presented relative to the control. Results are presented as mean values +/- SEM, N=3; *P < 0.05, one-tailed t-test, unequal variance.

**Supplemental Figure S2.**
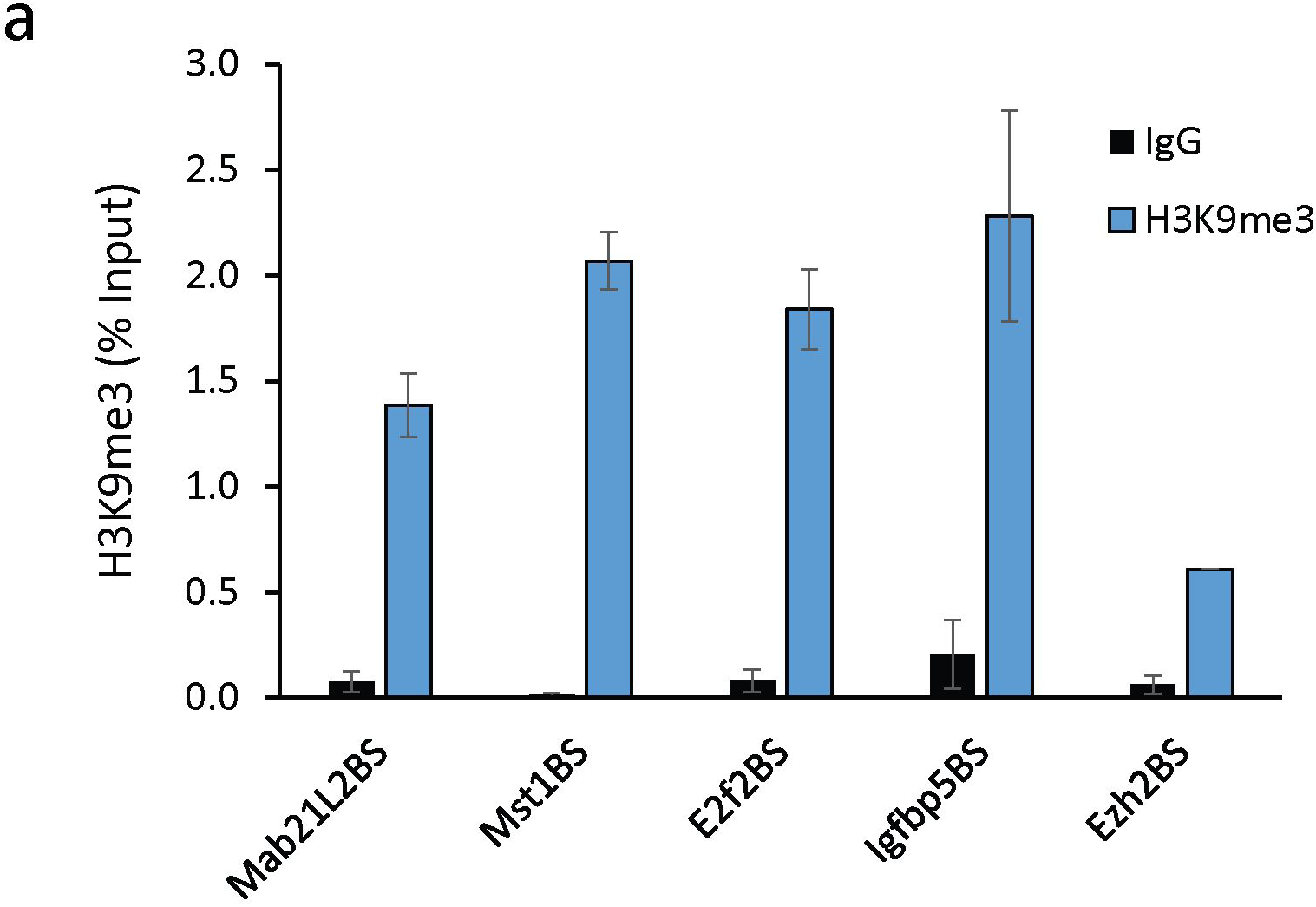
*Paupar*-KAP1-PAX6 bound sequences within the regulatory regions of the *Mab21L2*, *Mst1*, *E2f2*, *Igfbp5* and *Ezh2* genes are enriched in H3K9me3 modified chromatin. ChIP assays were performed in N2A cells using either histone H3K9me3 or anti-rabbit IgG control antibody. DNA fragments were amplified using qPCR. % input was calculated as 100*2^(Ct Input-Ct IP). Results are presented as mean values +/- SEM., N=4.

**Supplemental Figure S3.**
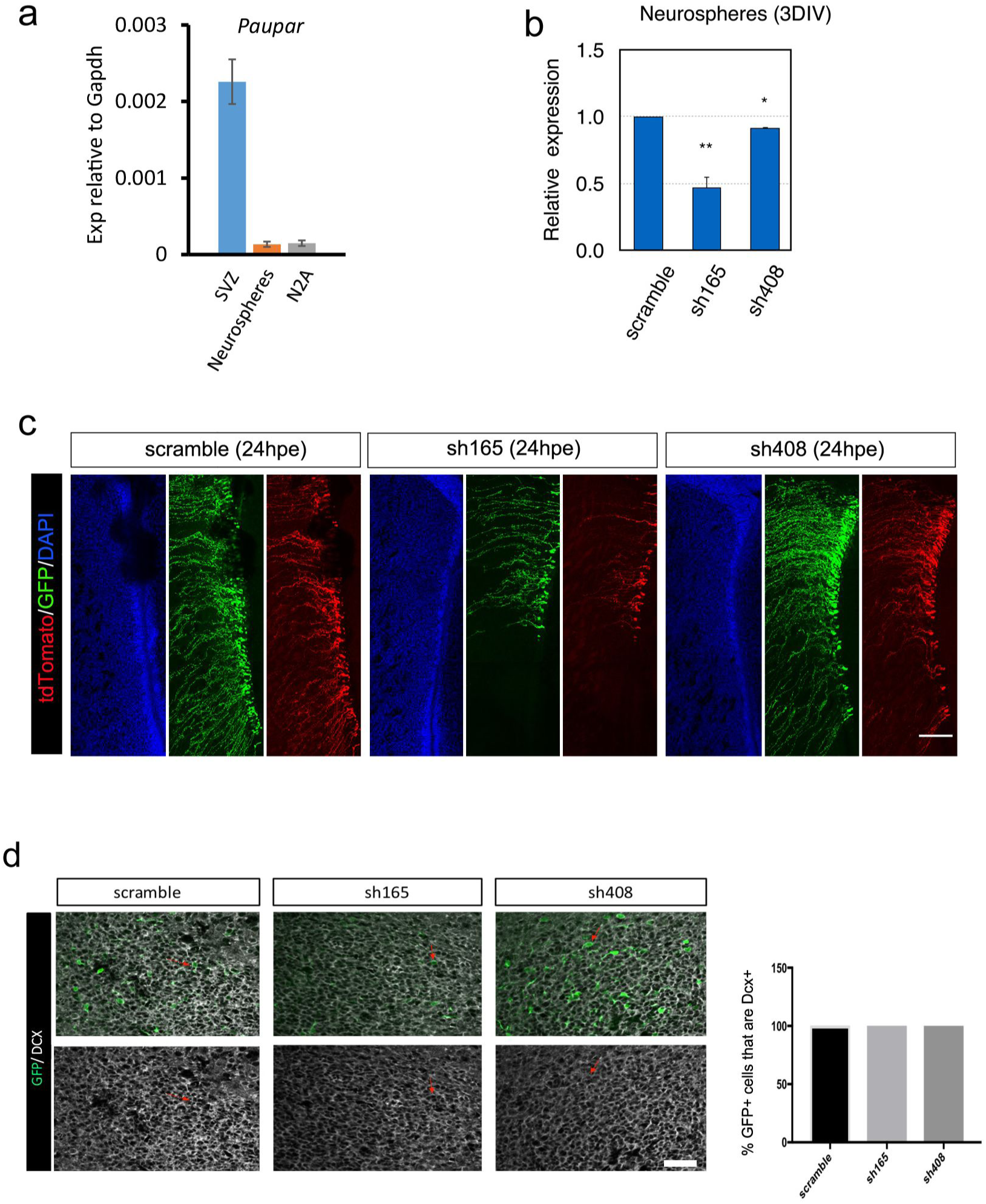
*Paupar* knockdown in SVZ. (a) *Paupar* transcript detected using RT-qPCR in the P4 SVZ and in tertiary neurospheres prepared from the P4 SVZ. (b) *Paupar* knockdown in tertiary neurospheres with sh165 and sh408. Neurosphere cultures were transfected with the indicated *Paupar* targeting shRNA expression vectors or a non-targeting control. *Paupar* expression was quantified using qRT-PCR three days later and normalised using *Gapdh*. The results are presented relative to the scrambled control (set at 1). (N≥4). (c) Immunostaining of GFP and mCherry in SVZ electroporated with shRNA and pCS-tdTomato at 24hpe. The concentration of the constructs were matched in order to minimize electroporation efficiency differences between scramble, sh165 and sh408. Fewer GFP+ cells were found in the SVZ after sh165 electroporation compared to scramble control electroporation. Co-electroporation with a construct expressing tdTomato driven by a different promoter (pCS), confirmed this was not due to the sh165 construct itself as fewer tdTomato+ cells were also observed at 24hpe. Knockdown with sh408 resulted in similar numbers of GFP+ electroporated cells compared to scramble controls at 24hpe. N=3. (d) GFP and DCX co-labelling in the OB 7dpe. Small red arrows show examples of co-labelled cells. There are no error bars because 100% of all GFP+ cells in the OB were DCX+. Data are shown as mean ± SEM and analysed by two-tailed Student t-tests. *p<0.05, **p<0.01, ***p<0.001. Scale bars represent 150 μm (c), and 30 μm (d).

**Supplemental Figure S4.**
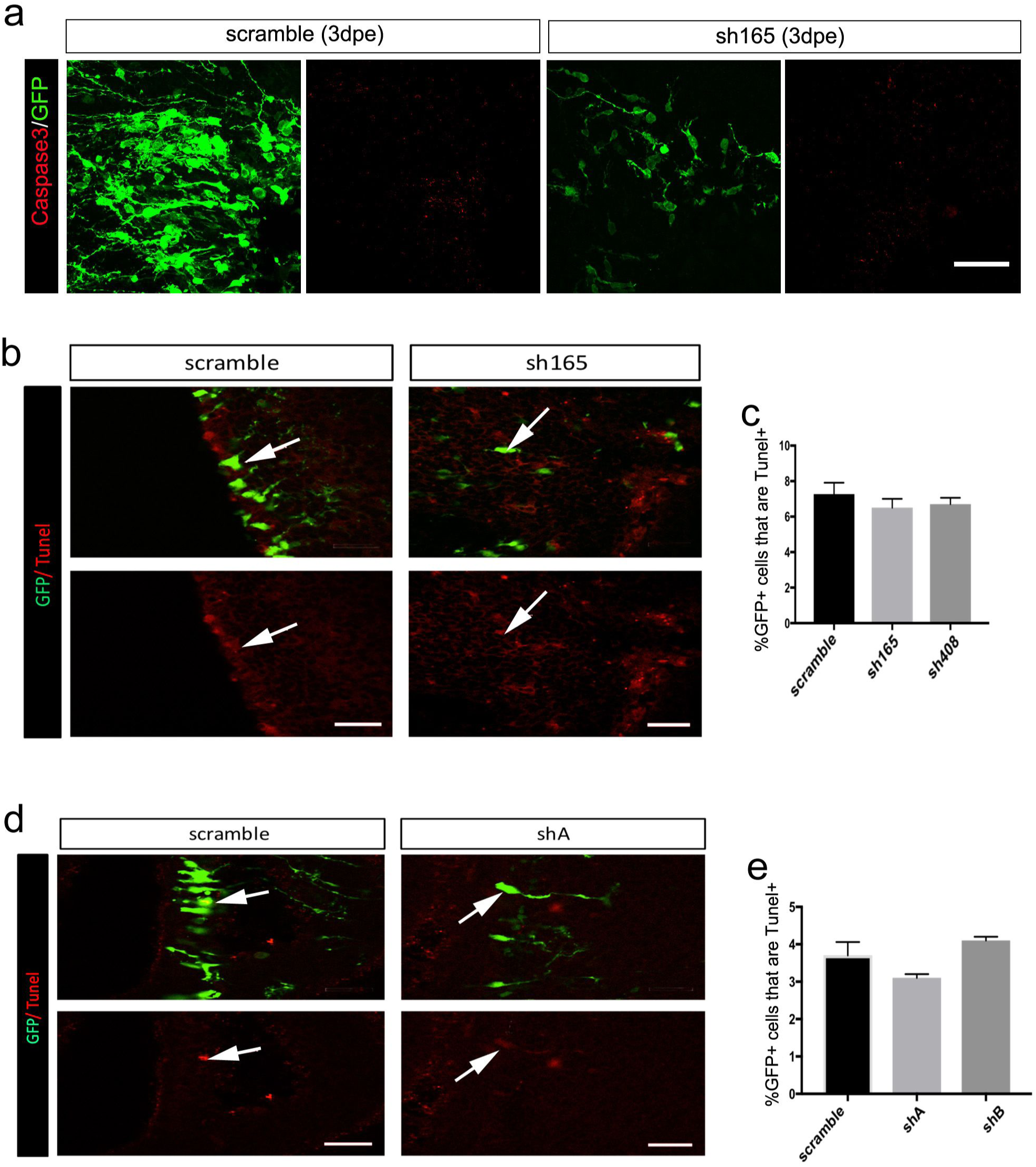
Cell death analysis after *Paupar* and Kap1 KD. (a) Immunostaining of GFP and CASPASE3 in SVZ at 3dpe. N=4. (b-c) Tunel assay and quantification in the SVZ after *Paupar* KD 3dpe. N=3. (d-e) Tunel assay and quantification in the SVZ after Kap1 KD 3dpe. N=3. Data are shown as mean ± SEM⍰and analysed by one-way ANOVA. Data are shown as mean ± SEM and analysed by two-tailed Student t-tests. Scale bars represent 50 μm (a), 30 μm (b,d).

